# A suite of pre-assembled, pET28b-based Golden Gate vectors for efficient protein engineering and expression

**DOI:** 10.1101/2025.01.13.632842

**Authors:** Deepika Gaur, Matthew L. Wohlever

**Affiliations:** Department of Cell Biology, University of Pittsburgh, Pittsburgh, PA 15261

**Author notes:** Correspondence to Matthew L. Wohlever.

**Keywords:** Protein engineering, protein expression, cloning, Golden Gate

## Abstract

Expression and purification of recombinant proteins in *E. coli* is a bedrock technique in biochemistry and molecular biology. Expression optimization requires testing different combinations of solubility tags, affinity purification techniques, and site-specific proteases. This optimization is laborious and time consuming as these features are spread across different vector series and require different cloning strategies with varying efficiencies. Modular cloning kits based on the Golden Gate system exist, but are overly complicated for many applications, such as undergraduate research or simple screening of protein purification features. An ideal solution is for a single gene synthesis or PCR product to be compatible with a large series of pre-assembled Golden Gate vectors containing a broad array of purification features at either the N or C-terminus. To our knowledge, no such system exists. To fulfill this unmet need, we Golden Gate domesticated the pET28b vector and developed a suite of 21 vectors with different combinations of purification tags, solubility domains, visualization/labeling tags, and protease sites. We also developed a completely scarless vector series with 9 different N-terminal tags. The system is modular, allowing users to easily customize the vectors with their preferred combinations of features. To allow for easy visual screening of cloned vectors, we optimized constitutive expression of the fluorescent protein mScarlet3 in the reverse strand, resulting in a red to white color change upon successful cloning. Testing with the model protein sfGFP shows the ease of visual screening, high efficiency of cloning, and robust protein expression. These vectors provide versatile, high-throughput solutions for protein engineering and functional studies in *E. coli*.

## Introduction

Recombinant protein expression and purification in *Escherichia coli* is a cornerstone technique in both biochemical research and the biotechnology industry^1,2^. This process begins with the cloning of the target gene into a plasmid that contains features to enhance protein expression and simplify purification, including affinity tags, solubility-enhancing domains, visualization/labeling tags, and protease cleavage sites for tag removal^3,4^. Achieving optimal expression is often challenging, especially for membrane proteins and other complex constructs^5–7^. Researchers frequently need to screen various combinations of these features as no single setup consistently yields the best results. Additionally, these features are typically spread across multiple vector systems, necessitating different and sometimes inefficient cloning strategies. Switching between these strategies to optimize expression and purification can be both time-consuming and resource intensive.

Golden Gate cloning has recently gained popularity due to its high efficiency and use of non-palindromic Type IIS restriction enzymes, which have separate recognition and cutting sites. Digestion with type IIS endonuclease creates custom overhangs that facilitate recombination-based cloning of compatible overhangs resulting in scarless cloning^8^. An added advantage of the Golden Gate vectors is the use of color based visual screen to differentiate between parental and cloned vectors^9–12^.

Modular cloning vector kits, like CIDAR and EcoFlex, are available to facilitate cloning using the Golden Gate method^13,14^. While these toolboxes provide great flexibility, the multicomponent kits are overly complicated for many applications, such as undergraduate research or simple screening of solubility and affinity purification tags. To our knowledge, no system exists wherein a researcher can use a single gene synthesis or PCR product that can be directly cloned into a large suite of pre-assembled Golden Gate vectors with varying combinations of tags and protease sites at both the N and C-terminus.

To address this issue, we domesticated the pET28b vector into the Golden Gate system. We then created three series of vectors containing N-terminal, C-terminal, or dual-tagged constructs with varying combinations of affinity purification tags, solubility enhancing domains, visualization/labeling tags, and protease cleavage sites. All vectors share compatible overhangs produced by digestion the Type IIS restriction enzyme Bsa1, thereby allowing a single gene synthesis to be directly cloned into 21 different pre-assembled vectors with the same cloning strategy. We also developed a completely scarless vector series with 9 different N-terminal tags. To allow for easy visual screening, we optimized constitutive expression of a red fluorescent protein in the parental vector, resulting in a red to white color change upon successful cloning. We demonstrate that these vectors have clear visual screening, high cloning efficiency, and robust protein expression. We envision this system will benefit researchers by significantly simplifying workflow and improving efficiency for recombinant protein expression and purification.

## Results

### Golden Gate domestication of pET28b

To domesticate the pET28b vector into the Golden Gate (GG) system we used the NcoI and XhoI restriction enzyme sites in pET28b to introduce outward-facing BsaI restriction sites and a fluorescent protein in the reverse strand for color-based screening of cloned vectors **(Figure 1A)**. The Golden Gate domesticated pET28b vector (pET28b-GG) retains all essential elements of the pET28b vector, ensuring robust expression under Isopropyl-β-D-Thiogalactopyranoside (IPTG) induction. To facilitate the insertion of tags at either terminus, our insert also contained NdeI and BamHI restriction enzyme sites. N-terminal tags can be cloned between NcoI and NdeI whereas C-terminal tags can be cloned between BamHI and XhoI **(Figure 1B-D)**.

**Figure 1:**
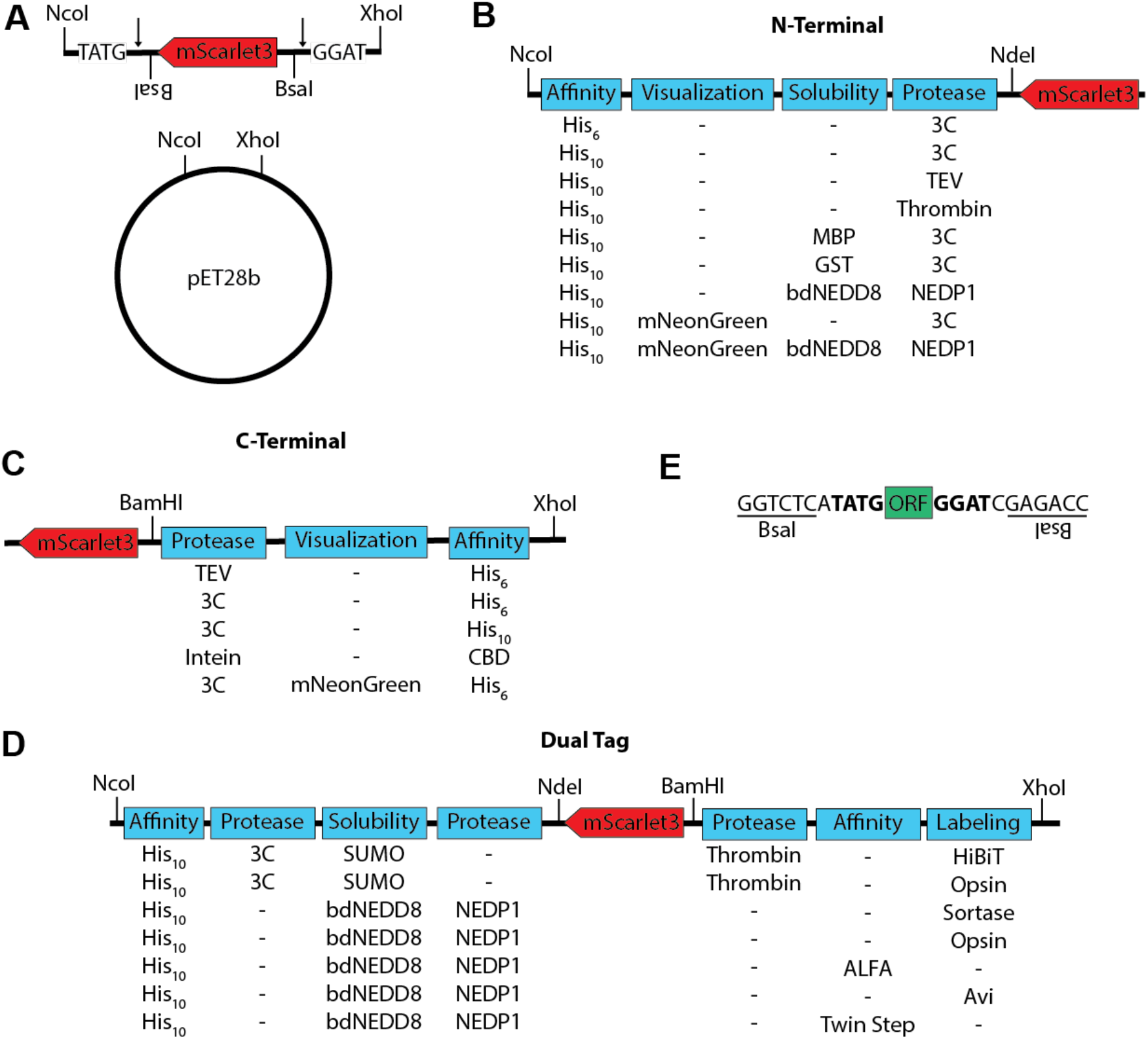
Development of the pET28b-GG vector series. **A)** The pET28b vector was Golden Gate domesticated by restriction cloning between NcoI and XhoI sites. The insert consists of mScarlet3 in the reverse strand with outward facing BsaI sites, which are marked by the arrows. **B)** Diagram of N-terminal vectors with affinity purification tags, visualization tags, solubility domains, and protease cleavage sites. Additional N-terminal features can be restriction cloned between NcoI and NdeI sites. **C)** Diagram of C-terminal vectors with protease sites, visualization tags, and affinity purification tags. Additional C-terminal features can be inserted between BamHI and XhoI. **D)** Diagram of Dual tagged vectors with affinity purification tags, protease sites, solubility domains, and labeling tags. Additional features can be inserted by restriction cloning with NcoI and NdeI for the N-terminus and BamHI and XhoI for the C-terminus. **E)** Diagram showing the 5’ and 3’ sequences that should be appended to the open reading frame (ORF) to facilitate cloning into pET28b-GG vectors. The BsaI sequences are inward facing with the compatible overhangs in bold.

After domesticating the vector, we expanded the versatility of the pET28b-GG by adding different affinity purification tags (His_6_, His_10_, CBD, MBP, GST, ALFA tag, and Twin Strep), solubility tags (MBP, SUMO, GST, and bdNEDD8), visualization/labeling tags (mNeonGreen, HiBiT, Sortase, Opsin, and Avi tag), and protease cleavage sites (3C, TEV, thrombin, NEDP1, and Intein)^15–23^. Many vectors contain multiple affinity purification tags, allowing for higher purity products via tandem affinity purification. This versatility supports a broad range of applications, such as protein purification, solubility enhancement, screening, and labeling which are critical for successful protein expression and downstream analysis.

In total, there are 21 different vectors that are fully compatible with the same insert **(Table 1)**. For simplicity, we divided these vectors into three different groups based on the location of the tags, which we refer to as N-terminal, C-terminal, and dual-tagged vectors **(Figure 1B-D)**. The system is expandable as new vectors with additional combinations of features can be added to the series via restriction cloning between NcoI and NdeI sites for N-terminal tags and BamHI and XhoI for C-terminal tags.

**Table 1:** List of pET28b-GG N-terminal, C-terminal, and dual tagged vectors. ORF is used to indicate where the desired open reading frame is in relation to the tags. All vectors listed in the table include a C-terminal glycine cloning scar.

| Plasmid # | Description |
| --- | --- |
| **N-terminal tag** |  |
| p344_DG | His<sub>6</sub>-3C-**ORF** |
| p345_DG | His<sub>10</sub>-3C-**ORF** |
| p346_DG | His<sub>10</sub>-TEV-**ORF** |
| p347_DG | His<sub>10</sub>-Thrombin-**ORF** |
| p348_DG | His<sub>10</sub>-MBP-3C-**ORF** |
| p349_DG | His<sub>10</sub>-GST-3C-**ORF** |
| p350_DG | His<sub>10</sub>-bdNEDD8-**ORF** |
| p351_DG | His<sub>10</sub>-mNeongreen-3C-**ORF** |
| p352_DG | His<sub>10</sub>-mNeongreen-bdNEDD8-**ORF** |
| **Dual Tag** |  |
| p361_DG | His<sub>10</sub>-3C-3xFLAG-SUMO-**ORF**- thrombin-HiBiT |
| p362_DG | His<sub>10</sub>-3C-3XFLAG-SUMO-**ORF**-thrombin-Opsin |
| p363_DG | His<sub>10</sub>-bdNEDD8-**ORF**-Sortase tag |
| p364_DG | His<sub>10</sub>-bdNEDD8-**ORF**-Opsin tag |
| p365_DG | His<sub>10</sub>-bdNEDD8-**ORF**-ALFA tag |
| p366_DG | His<sub>10</sub>- bdNEDD8-**ORF**-Avi tag |
| p367_DG | His<sub>10</sub>- bdNEDD8-**ORF**-2X Strep tag |
| **C-terminal tag** |  |
| p324_DG | **ORF**-TEV-His<sub>6</sub> |
| p368_DG | **ORF**-3C-His<sub>6</sub> |
| p369_DG | **ORF**-3C-His<sub>10</sub> |
| p370_DG | **ORF**-3C-mNeongreen-His<sub>6</sub> |
| p371_DG | **ORF**-Intein-Chitin Binding Domain |

All 21 vectors contain initiator methionine residues, stop codons, and outward-facing BsaI sites that generate unique 4-base overhangs upon digestion. The upstream overhang is TATG, which provides the initiator methionine. The downstream overhang is GGAT which encodes a glycine cloning scar. The cloning scar is necessary to allow the same gene synthesis product to be compatible with N-terminal, C-terminal, and dual-tagged vectors and we chose glycine because it is the smallest amino acid. To clone into the pET28-GG vectors, an insert should be flanked by GGTCTCA**TATG** at the 5’ end and **GGAT**CGAGACC at the 3’ end **(Figure 1E)**.

### Optimization of visual screening strategy to identify cloned vectors

A major advantage of different Golden Gate cloning kits is the incorporation of screening strategies to differentiate between undigested parental and cloned vectors. Golden Gate vectors often utilize a dropout strategy, where the parental vector encodes a fluorescent protein which results in the formation of colored colonies. Successful cloning results in the loss of the fluorescent protein and white colonies^9,24^.

This strategy has been successfully used with red fluorescent protein (RFP) in Golden Gate vectors intended for use in *S. cerevisiae*^24^. We attempted to replicate this strategy by cloning RFP into the reverse reading frame of the pET28b-GG vector, but we did not observe any color change **(Figure 2A)**. We hypothesized that the lac operator (LacO) sequence between the RFP open reading frame and Lac promoter was repressing transcription^25^. We therefore deleted the LacO sequence upstream of the RFP open reading frame. This resulted in formation of faint red color, which was still not distinct enough for clear visualization.

**Figure 2:**
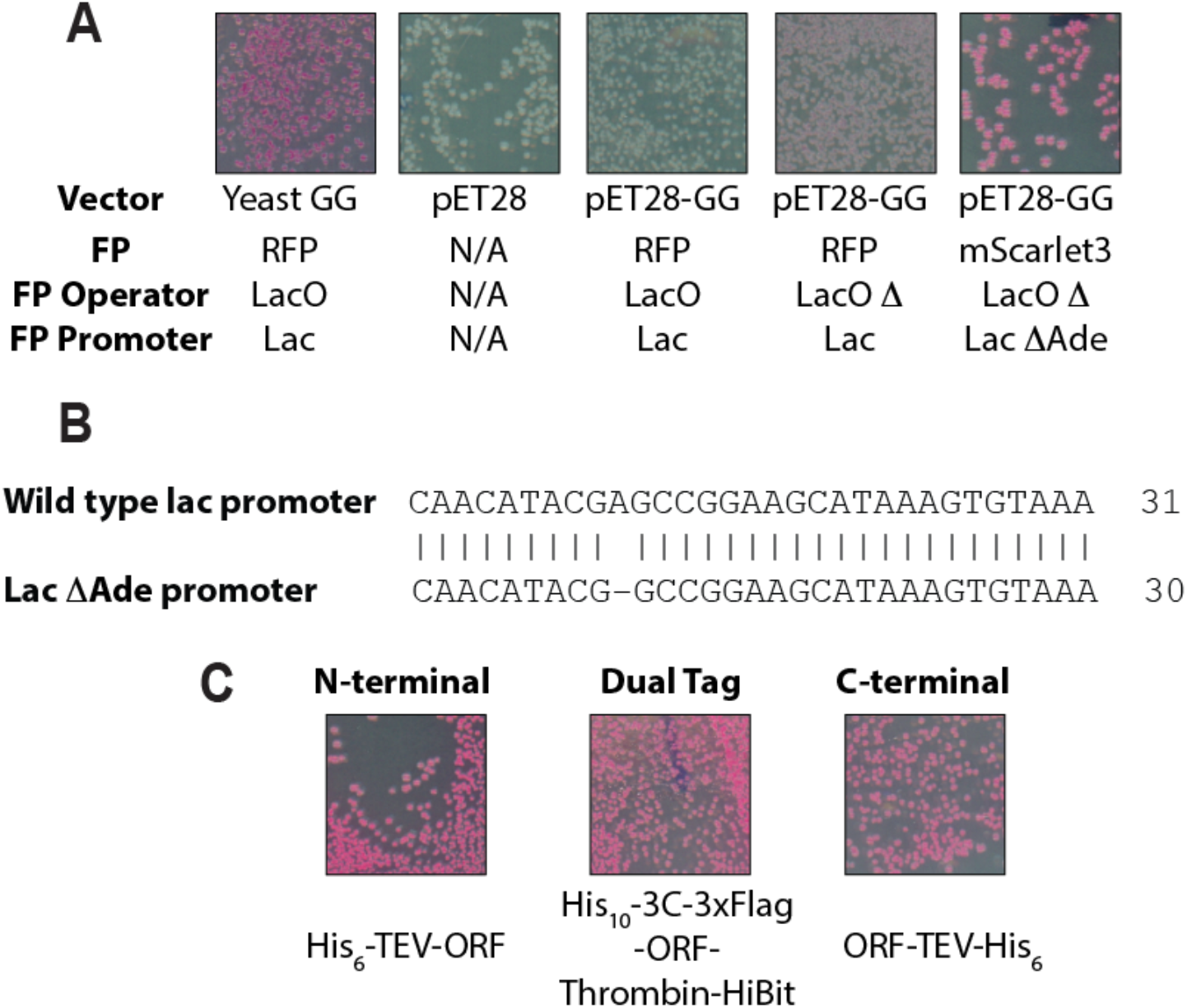
Optimization of color-based screening in the pET28b-GG vector. **A)** *E. coli* transformed with the pET28-GG vector exhibit robust red color when the fluorescent protein (FP) mScarlet3 is expressed from the Lac ΔAde promoter with the Lac Operator (LacO) deleted. **B)** The Lac ΔAde promoter has a single adenine deletion in the Lac promoter that give enhanced expression of mScarlet3. **C)** The robust visual screening is maintained in N-terminal, C-terminal, and dual tagged vectors.

A major difference between of *S. cerevisiae* (pRS series) and *E. coli* (pET28b) vectors is plasmid copy number. The pRS vectors utilize the pMB1 origin of replication, which has a copy number of 300-700 whereas pET vectors utilize the pBR322 origin, which has a copy number of 15-20^26^. To maintain as much similarity to the standard pET28b vector as possible, we chose not to change the origin of replication. We therefore decided to change our fluorescent protein to mScarlet3 which is 6.2 times brighter and has shorter maturation time compared to the RFP^27^.

After replacing RFP with mScarlet3, we noticed the presence of both faint and bright red colonies on the transformed plate. We selected the brightest colonies **(Figure 2A)** and performed whole plasmid sequencing. We discovered that the brightest colonies had a single base deletion in the lac promoter, which we refer to as the Lac ΔAde promoter **(Figure 2B)**. The combination of mScarlet3 and the Lac ΔAde promoter led to constitutive red color in parental pET28b-GG vectors. The red color was equally bright in the N-terminal, C-terminal, and dual-tagged vectors **(Figure 2C)**. We conclude that the combination of deleting the lac operator, the single base deletion in the lac promoter, and using the mScarlet3 reporter results in a robust, constitutive expression that is suitable for visual screening.

### Golden gate vectors have high cloning efficiency

To assess the cloning efficiencies of our vectors we cloned superfolder GFP (sfGFP) into a representative of each of the vector three classes. We chose His_10_-3C-ORF (p345_DG), ORF-TEV-His_6_ (p324_DG), and His_10_-3C-3xFlag-SUMO-ORF-thrombin-HiBiT (p361) as representatives of the N-terminal, C-terminal, and dual-tagged vectors, respectively. Our initial criterion for assessing the cloning efficiency is the presence of white colonies on LB plates after transformation. The red to white color shift indicates a positive clone that disrupted the expression of mScarlet3 in the parental vector. We observed 100% white colonies with each of the three vectors **(Figure 3A)**. Importantly, there were >100 colonies on each plate, indicating that the Golden Gate assembly reaction was highly efficient.

**Figure 3:**
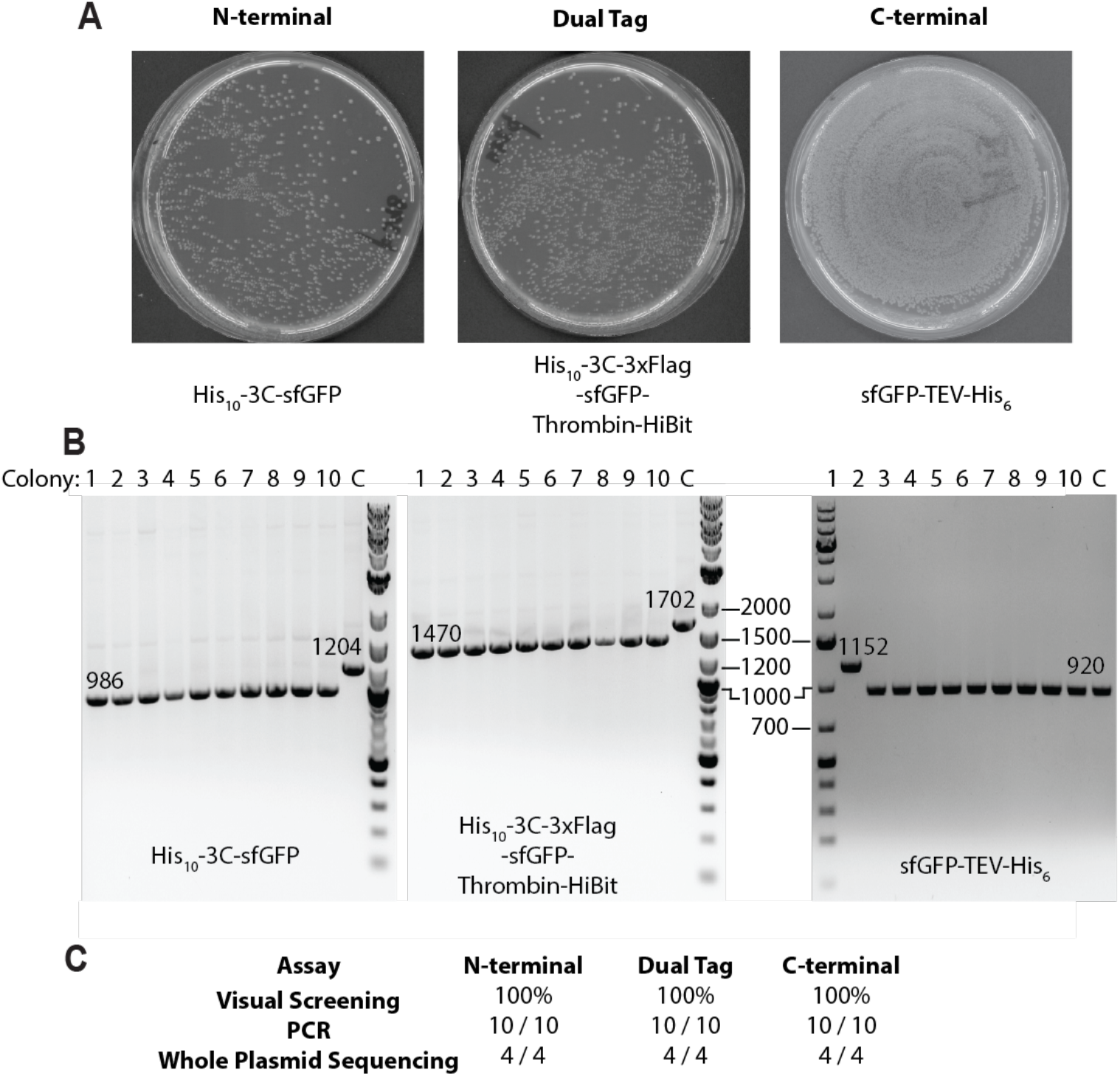
Cloning into pET28b-GG vectors is highly efficient. **A)** Golden Gate reactions were performed with representative vectors from the N-terminal, C-terminal, and dual tagged class of pET28-GG vectors. Imaging the plates 1 day after transformation shows 100% white colonies. **B)** Ten unique colonies from each of the above plates were cultured and miniprepped. The region between the T7 promoter and T7 terminator was PCR amplified and analyzed by agarose gel electrophoresis. The parental vector was used as a negative control (C). The expected size of the correct insert and the parental control are listed on the gel. All selected colonies have an insert of the correct size. Note that a different DNA ladder was used for the sfGFP-TEV-His_6_ gel. **C)** Summary of assays used to assess cloning efficacy in pET28b-GG vectors.

To determine whether the white colonies contained sfGFP we miniprepped DNA from 10 unique colonies from each transformation. We then used PCR to amplify the insert between the T7 promoter and terminator. The correctly cloned construct should have a smaller PCR product than the parental vector. Agarose gel analysis shows that 100% of the plasmids tested have an insert of the correct size. As a final control, we selected four plasmids from each vector series for whole plasmid sequencing. Again, 100% of vectors had the desired sequence **(Figure 3C)**. We conclude that the pET28b Golden Gate vectors have high cloning efficiency.

### pET28b GG vectors are suitable for protein overexpression and purification

To test the efficacy of the pET28b-GG vectors for protein expression and purification, we transformed the three representative plasmids expressing sfGFP into BL21 DE3 cells. Starter cultures were used to inoculate 1 L of terrific broth, which was incubated at 37° C until an OD_600_ of 0.6, at which point 0.3 mM IPTG was added to induce protein expression. After 16 h induction at 16° C, cells were pelleted, lysed, and purified by Ni-NTA chromatography. SDS PAGE analysis shows robust overexpression and purification of sfGFP with all three categories of vectors **(Figure 4)**. We conclude that the pET28 GG vectors are suitable for protein expression and purification.

**Figure 4:**
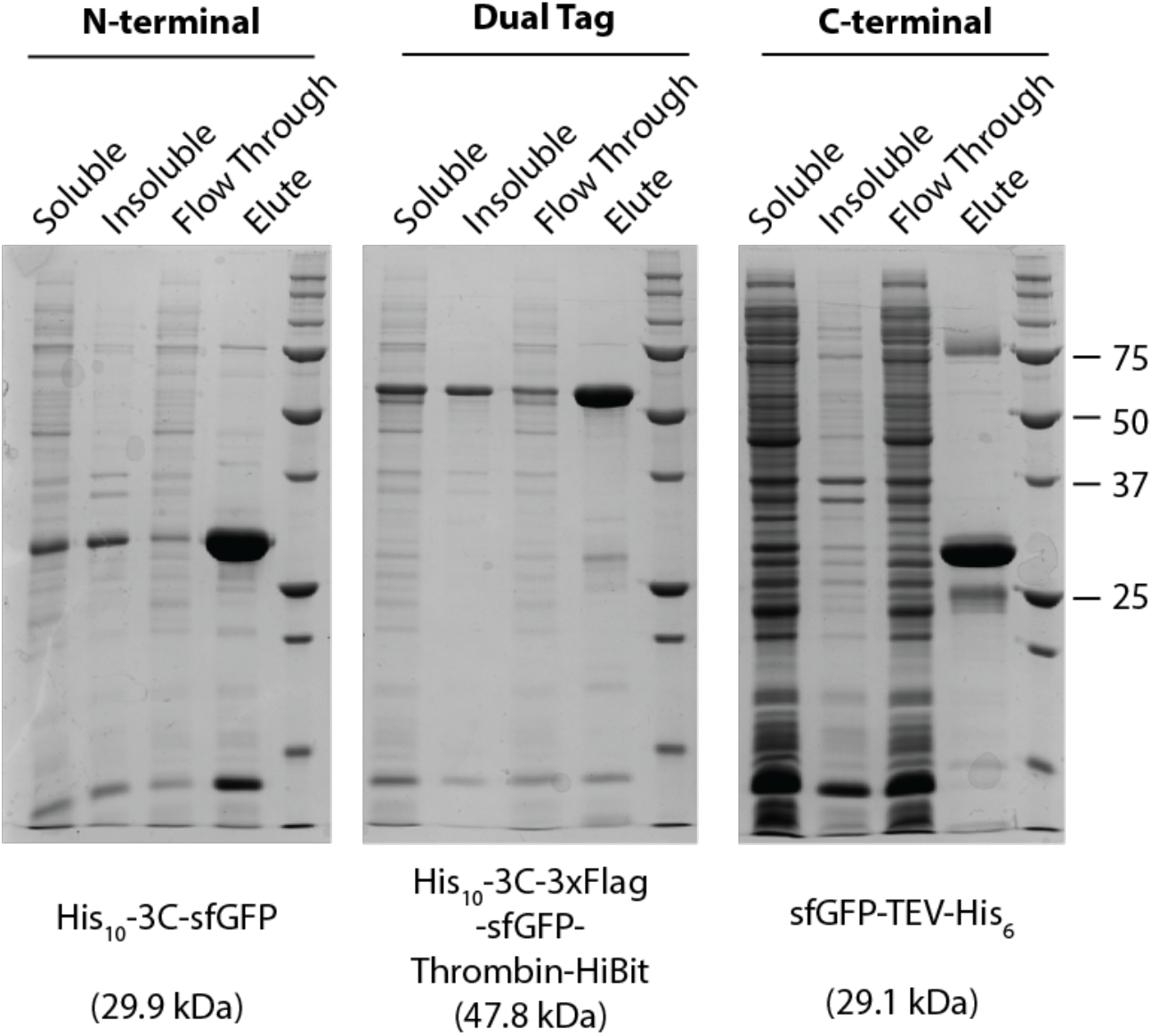
pET28b-GG vectors have robust protein overexpression. SDS-PAGE of the sfGFP purification from N-terminal, Dual, and C-terminal golden gate vectors.

### Design and Development of Scarless Vectors for N-Terminal Tagging

Our goal in designing the pET28b-GG vectors was to have a single gene synthesis product be compatible with an array of vectors with N-terminal, C-terminal, or dual terminal tags. This cross-compatibility necessitated a glycine scar at the C-terminus of all vectors. Although it is the smallest possible scar, the glycine scar may not be compatible with some downstream uses, such as structural biology.

To address this caveat, we generated a new series of vectors without any cloning scar, which we refer to as the scarless vectors. To eliminate the glycine scar, these vectors encode a stop codon. Therefore, the scarless vectors only have N-terminal tags and are not cross-compatible with C-terminal or dual tags.

The design of the scarless vectors is similar to the pET28b GG vectors with a Bsa1-mScarlet3-Bsa1 cassette. However, digestion with Bsa1 results in the generation of ATCC and TGAG overhang at the N-terminus and C-terminus respectively **(Figure 5A)**, which differs from the TATG and GGAT overhangs in the vector series described above. The scarless vectors encode the initiator methionine and stop codon so it is not necessary to include these in the insert. To clone into the scarless pET28-GG vectors, an insert should be flanked by GGTCTCA**ATCC** at the 5’ end and **TGAG**CGAGACC at the 3’ end **(Figure 5B)**. We generated a total of nine scarless vectors **(Table 2)**. The scarless vectors can be further customized by restriction cloning additional features between the NcoI and BamHI sites **(Figure 5C)**. Similar to the pET28b-GG vectors, there is robust expression of mScarlet3 which allows for easy visual screening of colonies **(Figure 5D)**.

**Table 2:**
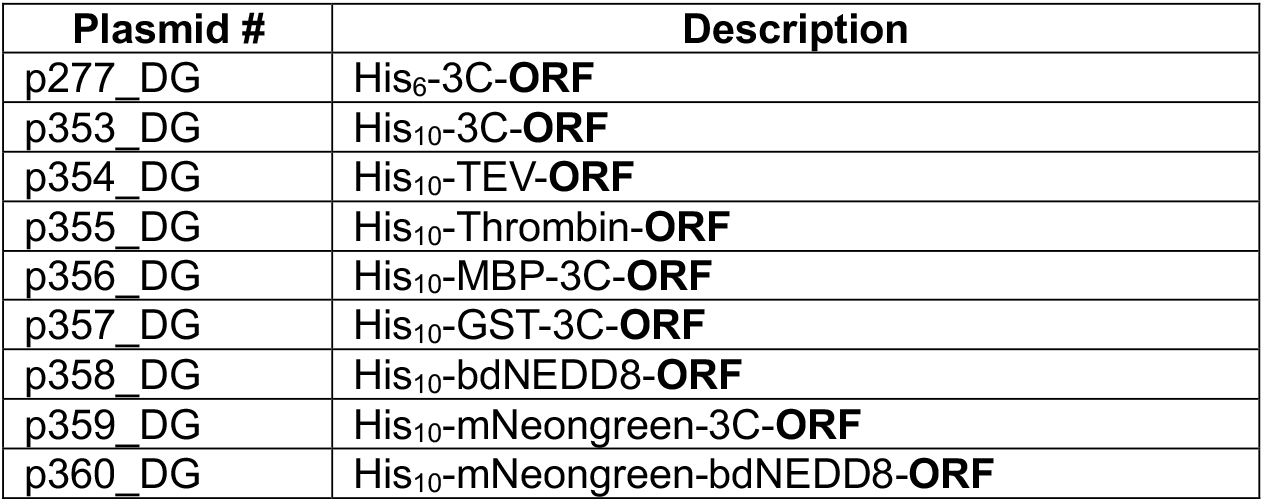
Scarless N-terminal vectors. ORF is used to indicate where the desired open reading frame is in relation to the tags. All vectors listed in the table are scarless.

**Figure 5:**
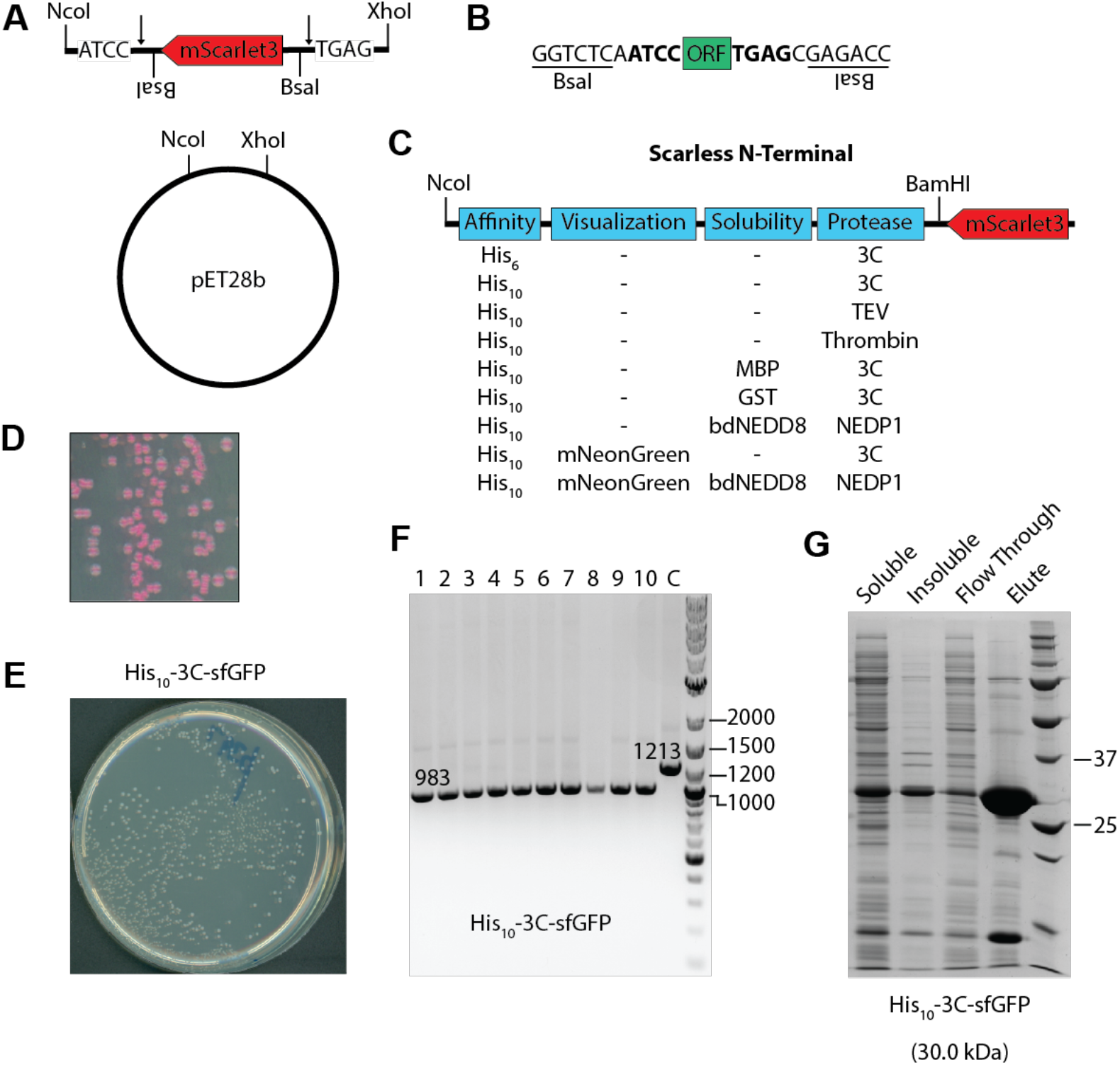
Design and validation of Scarless N-terminal pET28b-GG vectors. **A)** Schematic of the scarless N-terminal vectors. The pET28b vector was Golden Gate domesticated by restriction cloning between NcoI and XhoI sites. The insert consists of mScarlet3 in the reverse strand with outward facing BsaI sites. **B)** Diagram showing the 5’ and 3’ sequences that should be appended to the open reading frame (ORF) to facilitate cloning into the scarless pET28b-GG vectors. The BsaI sequences are inward facing with the compatible overhangs in bold. **C)** Diagram of N-terminal vectors with affinity purification tags, visualization tags, solubility domains, and protease cleavage sites. Additional N-terminal features can be restriction cloned between NcoI and NdeI sites. **D)** The robust visual screening is maintained in scarless N-terminal vectors. **E)** Golden Gate reactions were performed with a representative scarless N-terminal pET28-GG vector. Imaging the plates 1 day after transformation shows 100% white colonies. **F)** Ten unique colonies from the scarless N-terminal vector were cultured and miniprepped. The region between the T7 promoter and T7 terminator was PCR amplified and analyzed by agarose gel electrophoresis. The parental vector was used as a negative control (C). All selected colonies have an insert of the correct size. **G)** SDS-PAGE of the His_10_-3C-sfGFP purification from Scarless N-terminal vector.

To validate the cloning efficacy of the scarless vectors, we cloned sfGFP into the scarless His_10_-3C-ORF vector (p353_DG). Upon transformation, we detected 100% white colonies **(Figure 5E)**. We selected 10 colonies for PCR amplification and observed that all 10 colonies gave PCR products of the expected size **(Figure 5F)**. We then selected for plasmids for whole plasmid sequencing, which again confirmed that 100% of the colonies had the correct sequence.

To test the efficacy of the scarless vectors for protein expression and purification, we transformed the scarless His_10_-3C-sfGFP vector into BL21 DE3 cells and used a starter culture to inoculate 1 L of terrific broth. After overnight IPTG induction at 16° C, cells were pelleted, lysed, and purified by Ni-NTA chromatography. SDS PAGE analysis shows robust overexpression and purification of sfGFP from the scarless vector **(Figure 5G)**. We conclude that the scarless vectors have robust cloning and protein expression.

## Discussion

A major bottleneck in recombinant protein production is screening through various combinations of features to optimize protein expression and purification^4^. This laborious procedure often involves cloning the target gene into diverse vectors via different cloning strategies with variable efficiency. While modular cloning strategies such as CIDAR and EcoFlex offer modular feature assembly and efficient Golden Gate cloning, they are excessively complex for routine applications or educational settings.

Here we describe the creation and validation of a diverse range of Golden Gate-compatible pET28b vectors that facilitate rapid screening and optimization of recombinant protein expression in *E. coli*. These newly developed vectors allow a single PCR product to be cloned into a collection of 21 pre-assembled Golden Gate vectors with different combinations of affinity purification tags, solubility enhancing domains, labeling/visualization tags, and protease sites at both the N- and C-terminus. Because cross-compatibility with N- and C-terminal tags necessitates a glycine cloning scar, we also developed a set of nine scarless pET28-GG vectors with N-terminal tags. The Golden Gate domestication of the pET28b vector builds upon its well-established role in protein expression while introducing improvements such as incorporation of tandem affinity purification tags and solubility enhancers which enable higher purity and yield of target proteins. The system is expandable, allowing for the creation of new vectors with additional feature combinations via restriction cloning.

A key enhancement of our pET28b-GG vectors is the enhanced visual screening achieved through constitutive expression of the bright fluorescent protein mScarlet3. Test results with the model protein sfGFP demonstrate clear visual screening, high cloning efficiency, and robust protein expression in all classes of pET28b-GG vectors. One limitation is that we did not modify key vector features such as the copy number, promoter, or other regulator elements for expression of the target protein. Future work will focus on applying the strategy described here to other workhorse vectors in recombinant protein expression.

In conclusion, the pET28b-GG vector series provides a reliable, efficient, and user-friendly solution for recombinant protein expression and purification. The modular design and compatibility with a single cloning strategy streamlines workflows and make it an excellent tool for both experienced researchers and educational laboratories.

## Materials and Methods

### Golden Gate domestication of pET28b vector

The empty pET28b vector was a kind gift of the Keenan lab^28^. To Golden Gate domesticate the pET28b vector, a DNA construct consisting of mScarlet3 flanked by outward facing BsaI sites, a 5’ NcoI site, and 3’ XhoI site was generated by gene synthesis. The gene synthesis product was restriction cloned into pET28b. N-terminal features were either PCR amplified from existing vectors in the lab or generated by gene synthesis and restriction cloned in via NcoI and NdeI restriction sites. C-terminal tags were likewise generated by PCR or gene synthesis and restriction cloned between BamHI and XhoI. For the scarless N-terminal vectors, N-terminal features were generated by PCR or gene synthesis and restriction cloned between NcoI and BamHI sites. All plasmids were validated by whole plasmid sequencing.

### sfGFP cloning in Golden Gate vectors

To clone sfGFP into the pET28b-GG vectors, PCR primers were designed to amplify residues 2-239 of sfGFP. The primers also contained the appropriate 5’ and 3’ flanking sequences indicated in **Figures 1 & 5**. PCR products were run on agarose gel and purified by gel extraction (Qiagen).

Golden Gate assembly was performed as described earlier^24^. Briefly, 100 ng of the appropriate pET28b GG vector was mixed with sufficient insert from PCR amplification to achieve a 1:2 molar ratio of vector to insert. The reaction volume was brought up to a final volume of 15 μL with T4 DNA ligase buffer (NEB, B0202S), 200 U of T4 DNA ligase (NEB, M0202L), 20 U of Bsa1 (NEB, R3733L), and deionized water purified by reverse osmosis. All the reaction components are mixed in the ice to avoid degradation of ATP in the T4 DNA ligase buffer. The reaction was incubated on a thermocycler with 25 cycles of 37° C for 3 minutes and 16° C for 4 minutes. After the 25 cycles, the sample was incubated at 50° C for 5 minutes and 80° C for 5 minutes. Following thermocycling, 3 μL of the reaction was transformed into the DH5α cells and plated on the LB plates containing kanamycin and incubated overnight at 37° C.

### PCR verification of cloning efficiency

To validate cloning efficiency, 10 white colonies from each plate were used to inoculate 5 mL cultures of luria broth (LB) and grown to saturation. Plasmids were isolated with QIAGEN’S QIAprep spin miniprep kit. The insert was amplified with primers the bind to the T7 promoter (TAATACGACTCACTATAGGG) and T7 terminator (GCTAGTTATTGCTCAGCGG). The parental pET28b GG vector was used as a negative control. PCR consisted of a final concentration of 2 ng/ μL of plasmid, 10 μM of the forward and reverse primer, 200 μM dNTPs, and 1 U of Phusion DNA polymerase (Thermo Scientific). Thermocycling was performed as follows: 98° C for 30 seconds, then 24 cycles at 98° C for 30 seconds, 60° C for 30 seconds, 72° C for 1 minute, and finally for 1 minute at 72° C. Samples were analyzed on 1% agarose gel and then 4 colonies were selected for whole plasmid sequencing using service from Plasmidsauras.

### Protein expression and purification

Protein expression and purification were carried out as previously described^29^. Vectors encoding sfGFP were transformed into BL21 DE3 cells with pRIPL plasmid for rare codons. Individual colonies were selected and used to start a 5 mL LB culture. This culture was grown at 37° C for 6 hours and then used to inoculate a 15 mL of terrific broth (TB). The TB culture was grown at 37° C until saturation and then 10 mL of this culture was used to inoculate 1 L of TB. The 1 L culture was grown at 37° C until an OD_600_ of 0.6. Protein expression was then induced by the addition of isopropyl β-D-1 thiogalactopyranoside (IPTG) at a final concentration of 300 μM.

After addition of IPTG, the temperature was dropped to 16° C and the cells were grown for an additional 16 h. Cells were pelleted by centrifugation and resuspended in the Lysis Buffer (20 mM Tris pH 7.5, 200 mM potassium acetate, 20 mM imidazole, 1 mM DTT, 0.01 mM EDTA and 10% glycerol) supplemented with 0.05 mg/mL Lysozyme and 1 mM phenylmethylsulfonyl fluoride (PMSF).

For purification, cells were lysed by sonication and supplemented with 250 U of universal nuclease (pierce). The insoluble fraction was separated by centrifugation at 20,000 x g for 30 minutes. The supernatant was then incubated with 1 mL of packed Ni-NTA resin (Qiagen) and incubated in the cold room with gentle agitation for 30 minutes. The Ni-NTA resin was pelleted by centrifugation 5,000 x g for 5 minutes, the supernatant was decanted, the resin was resuspended in 5 mL of Lysis Buffer and loaded onto a gravity column where it was washed with 15 column volumes of Lysis Buffer. The resin was further washed with 5 column volumes of Wash Buffer (20 mM Tris pH 7.5, 200 mM potassium acetate, 30 mM imidazole, 1 mM DTT, 0.01 mM EDTA and 10% glycerol). Protein was eluted with 5 column volumes of Elution Buffer (20 mM Tris pH 7.5, 200 mM potassium acetate, 250 mM imidazole, 1 mM DTT, 0.01 mM EDTA and 10% glycerol). Protein purification was then analyzed by SDS PAGE on a 15% acrylamide gel.

## Abbreviations and symbols

3C: Human rhinovirus 3C protease
bdNEDD8: NEDD8 from *Brachypodium distachyon*
CBD: Chitin Binding Domain
GST: Glutathione S-transferase
MBP: Maltose binding protein
TEV: Tobacco Etch Virus

## Acknowledgements

The authors wish to thank members of the Wohlever lab for helpful discussions and feedback on the project.

## Author Contributions

Conceptualization: DG, MLW; Data curation: DG, MLW; Formal analysis: DG, MLW; Funding acquisition: MLW; Investigation: DG; Methodology: DG; Project administration: MLW; Resources: DG, MLW; Software: Not applicable; Supervision: MLW; Validation: DG; Visualization: DG, MLW; Writing – original draft: DG; Writing – review & editing: DG, MLW;

